# Invisible flashes bias spatial coding in the auditory system

**DOI:** 10.64898/2026.02.11.705146

**Authors:** Patrycja Delong, Claire Pleche, Uta Noppeney

## Abstract

Information integration is widely considered fundamental for human consciousness. Combining spatial ventriloquism and dynamic continuous flash suppression, this psychophysics-EEG study investigated how invisible flashes influence spatial perception of sounds. We show that invisible flashes, even when mislocalized by observers, bias observers’ perceived sound location. EEG multivariate pattern analyses unravelled that the locations of invisible flashes are encoded in early (150-250ms) neural activity. Crucially, even though this visuospatial information rapidly dissipates for invisible mislocalized flashes, being no longer decodable from neural activity after 300ms, it progressively biases neural encoding of sound location (300-600ms), thereby ultimately inducing a perceptual ventriloquist illusion.

Our results demonstrate that subjective awareness is not a prerequisite for integrating information across the senses. Spatial information that dissipates from the visual system and evades awareness biases neural encoding and perception of sound location. These results pose challenges for leading theories of consciousness as a unified ‘all-or-none’ phenomenon.

## Introduction

Information integration is widely regarded as a hallmark of human consciousness (Baars, 2005; Baars & Owen, 2002; Dehaene & Changeux, 2011; Lamme, 2006; Mudrik et al., 2014; Tononi, 2004; Tononi et al., 2016). Notably, the Global Workspace Theory assumes that nonconscious processing is confined to local neural circuitries, whereas conscious information enters a global workspace and is broadcast across distant brain regions via long-range connectivity. Because multisensory perception inherently relies on communication between sensory systems, it is ideally suited to test key predictions of theories of consciousness.

Recent psychophysics studies have shown that supraliminal stimuli from one sensory modality can boost subliminal stimuli from another modality into perceptual awareness (Adam & Noppeney, 2014; Aller et al., 2015; Alsius & Munhall, 2013; Cox & Hong, 2015; Lunghi et al., 2010, 2017; Lunghi & Alais, 2015; Ngo & Spence, 2010; Olivers & Van der Burg, 2008; Salomon et al., 2015, 2017; Zhou et al., 2010). These influences from supraliminal to subliminal processing are not surprising, because the neural activity associated with processing of conscious signals can enter the global workspace and thereby influence local processing of subliminal signals in other sensory modalities.

Initial psychophysics work has also revealed unconscious cross-modal associative learning (Scott et al., 2018) congruency priming (Faivre et al., 2014) and influences of unconscious visual signals on conscious sound perception(Delong et al., 2018; Delong & Noppeney, 2021). Most prominently, an invisible flash that was presented displaced from a concurrent sound induced a ventriloquist illusion, i.e. a shift or bias in observers’ perceived sound location towards the flash location (Delong et al., 2018; Delong & Noppeney, 2021). Computationally these crossmodal biases arise, because the brain forms an audiovisual spatial estimate by integrating auditory and visual spatial information weighted by their relative reliabilities (Aller & Noppeney, 2019; Meijer et al., 2019; Rohe & Noppeney, 2015a, 2015b). The impact of ‘unaware’ visual information on conscious sound perception poses challenges for the Global Workspace Theory in which consciousness arises as a unified ‘all-or-none’ phenomenon via ignition. It raises the critical question of how spatial information about a flash that evades observers’ subjective awareness in the visual system interacts with conscious spatial representations in the auditory system.

This psychophysics and four-day EEG study investigated how flashes that evade observers’ visual awareness influence the neural encoding and conscious perception of sounds. Combining dynamic Continuous Flash Suppression (dCFS) (Maruya et al., 2008; Tsuchiya & Koch, 2005) with spatial ventriloquism we presented observers on each trial with an auditory burst of white noise and a synchronous flash at congruent or incongruent locations along the azimuth under dCFS (Maruya et al., 2008). On each trial, observers rated the flash’s visibility and located the flash and the sound. Using multivariate decoding models, we temporally resolved the neural encoding of spatial information about visible and invisible flashes. Next, we investigated whether and how these visible and invisible flashes influence the neural encoding and perception of sound location. Our initial analyses focus on subjective awareness (i.e. visibility ratings), because they are closest to observers’ phenomenal experience and thereby key for assessing the Global Workspace Theory . Yet, visibility ratings are susceptible to criterion shifts (Dehaene & Changeux, 2011). We therefore also assess how invisible flashes bias the neural encoding and perception of sound location (i.e. neural and behavioural ventriloquist effect) depending on whether observers localized the invisible flash correctly or incorrectly. Our main results show that spatial information about invisible and incorrectly localized flashes rapidly dissipates from the visual system, being no longer decodable after 330 ms. Yet, this spatial information can progressively bias the neural encoding of sound location, thereby eventually leading to observers’ experience of a ventriloquist illusion.

## Results

72 participants were included in the analysis of the one-day psychophysics experiment (see inclusion criteria in Methods section). 18 participants completed the four days of testing for the subsequent EEG experiment. Both psychophysics and EEG experiments presented observers with an auditory burst of white noise emanating from one of three potential locations: left, centre or right. In synchrony with the sound, one eye was presented with i. no flash or a brief flash in the ii. left or iii. right hemifield under dynamic continuous flash suppression (dCFS) to the other eye (Maruya et al., 2008). Hence, the 3 x 3 factorial design manipulating i. ‘flash’ (left, right, absent) and ii. ‘sound location’ (left, central, right) (Figure 1A). In addition, the flash was presented either in the upper or lower hemifield. On each trial observers reported their perceived i. sound location (left, middle, right), ii. flash location (up vs. down) and rated iii. the flash’s visibility (clear image, almost clear image, weak glimpse, not seen) with the order of questions counterbalanced across participants. Visual and auditory spatial classification tasks thus focussed on orthogonal spatial dimensions to minimize response interference across tasks.

**Figure 1.**
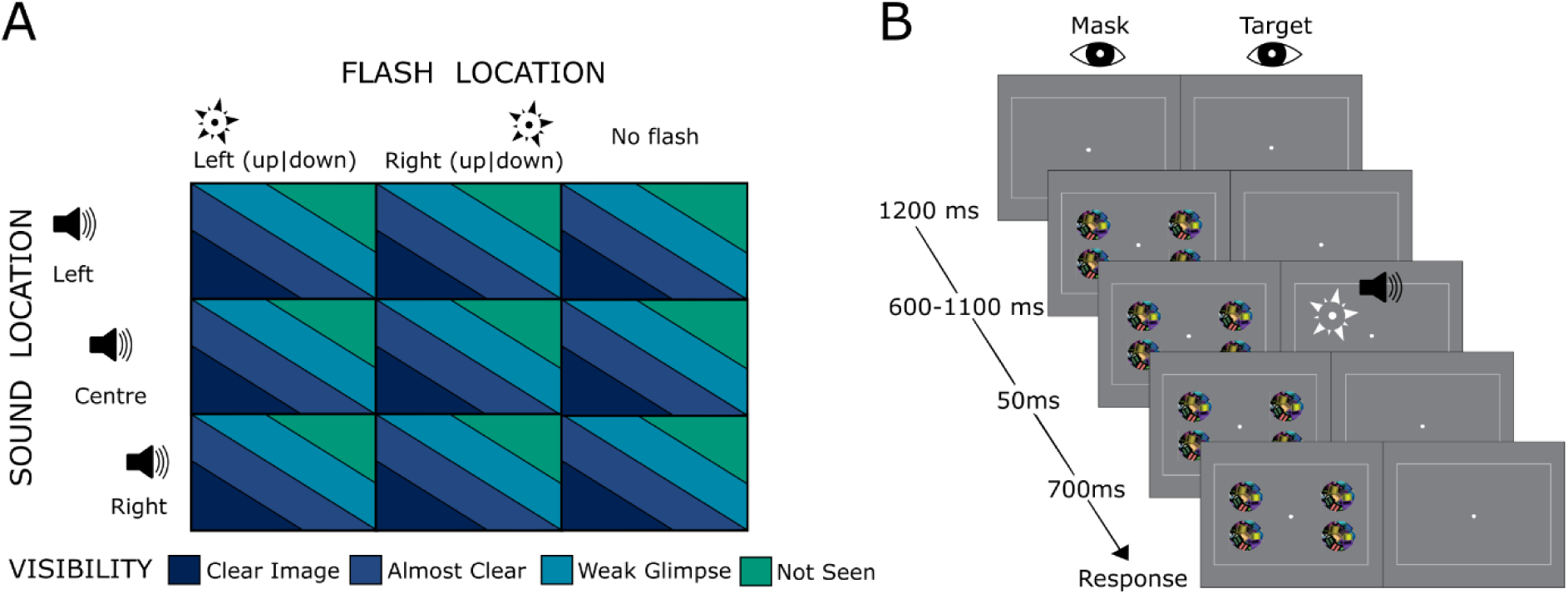
Experimental paradigm and procedure. A. Experimental design: 3 × 3 factorial design with the factors: (1) Flash: left (up|down), right (up|down), no flash; (2) Sound location: left, centre, right. The trials were categorized according to participants’ subjective visibility: Clear Image, Almost Clear Image, Weak Glimpse, Not Seen. B. Timecourse of one example trial.

dCFS obliterated subjective visual awareness and accurate flash localization only on a fraction of trials. This allowed us to compare the impact of physically identical flashes on observers’ perceived/reported (i.e. behavioural ventriloquist effect) and neurally encoded (i.e. neural ventriloquist effect) sound location depending on whether the flash was i. visible or invisible and ii. localized correctly or incorrectly by observers. Moreover, we finally assessed how the neural ventriloquist effect for visible and invisible flashes is related to observers’ experience of a perceptual ventriloquist illusion.

### Behavioural results

Sound localization accuracy of 65% ± 3.2 correct on the unisensory auditory trials indicated that observers were able to locate the sound. Likewise, their visibility ratings showed that dCFS was effective in manipulating their subjective, awareness. About 52% ± 3.7 of the trials were rated as invisible. Furthermore, observers’ accuracy on flash localization decayed consistently with their subjective visibility ratings from above 90% accuracy for visible trials to 54% accuracy for invisible trials. These results confirm that observers were attentive to the flash localization task and their flash visibility ratings adequately assessed whether they perceived the flash. To ensure reliable parameter estimation, we reduced the four visibility levels to two visibility levels: 1. Visible included: ‘clear image’, ‘almost clear image’ or ‘weak glimpse’ and 2. Invisible = ‘not seen’. We thereby included trials as invisible, only when they obtained the most stringent ‘invisible’ ratings, though even on those subjectively invisible trials observers’ flash spatial discrimination accuracy was above chance performance (right-tailed one sample t-test: t(17) = 3.576, p = 0.001).

#### The behavioural ventriloquist effect, depending on flash visibility and localizability

In the psychophysics and the EEG experiments, we first assessed how the flash location biases observers’ perceived sound location as quantified by the ventriloquist effect depending on flash visibility and observer’s localization accuracy. We computed the behavioural spatial ventriloquist effect as the perceived sound location for ‘visual right’ minus ‘visual left’ (averaged across trials) separately for visible and invisible trials (see Figure 2A left). Consistent with previous research, we observed a significant ventriloquist effect (VE) for both visible and invisible flashes (see Table 1, Figure 2A right, (Delong et al., 2018). Further, spatial ventriloquism was significantly greater for visible relative to invisible flashes as indicated by a paired t-test (psychophysics experiment: t(71) = 10.889, p < 0.001; EEG experiment: t(17) = 6.031, p < 0.001). Because the ventriloquist effect was computed averaged across trials, this difference in ‘VE magnitude’ mainly reflects the fact that the proportion of trials on which observers experienced a ventriloquist illusion was significantly greater for visible than invisible flashes (percentage of trials with behavioural ventriloquist effect for visible trials). In the EEG experiments, we observed a ventriloquist effect for sounds at the centre location on 59% ± 7.3 of the visible trials and on 31% ± 7.2 of the invisible trials (two-tailed paired t-test comparing visible and invisible trials: t(17) = 4.83, p < 0.001).

**Figure 2.**
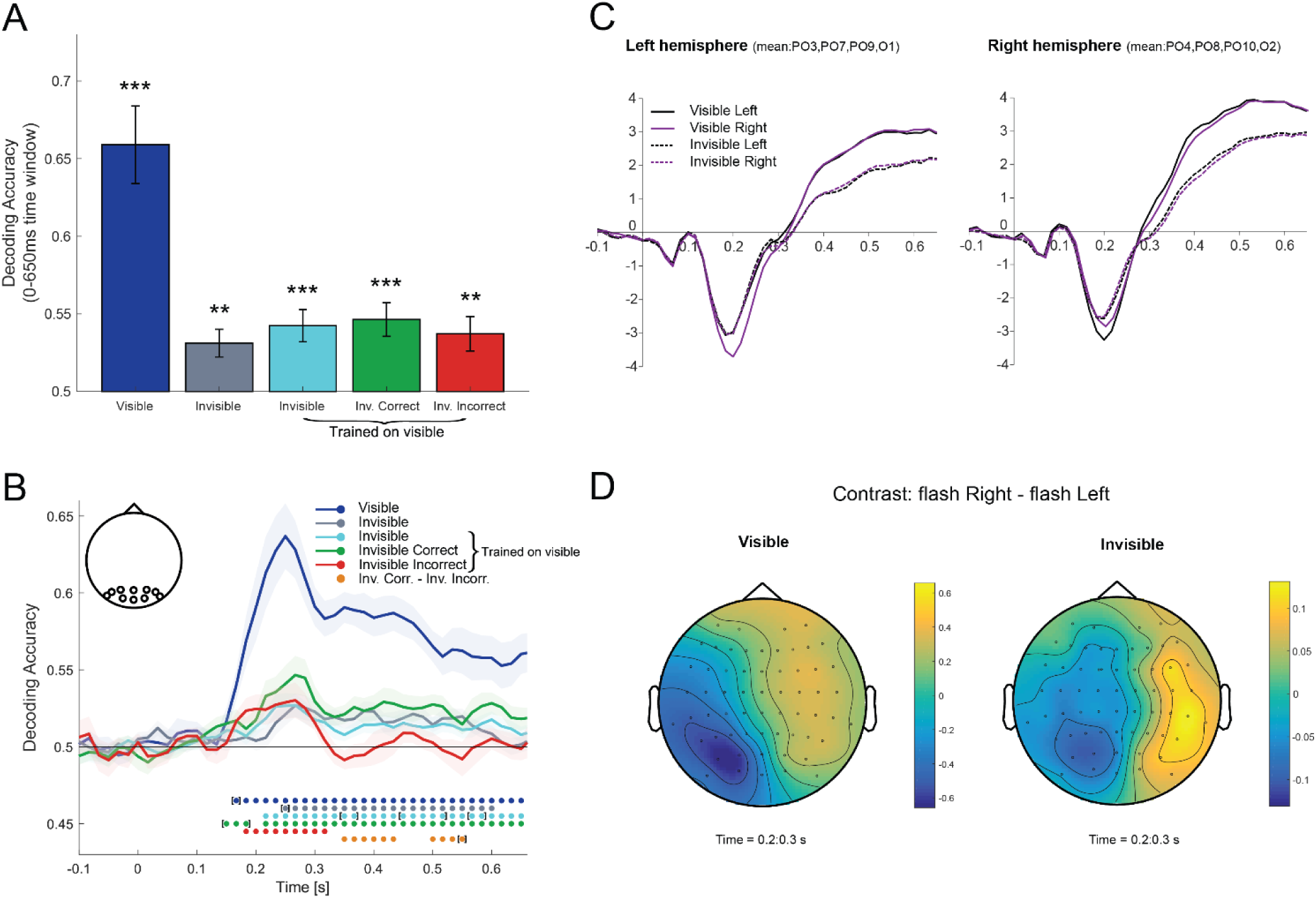
Visual-spatial representations. A. Bar plots showing EEG decoding accuracy for Left vs Right flash location (across subjects mean ± SEM). B. Time course of classifier’s accuracy. Dots without brackets indicate the time points with classifier’s performance significantly better than chance at p < 0.05, and dots with brackets indicate the time points with classifier’s performance marginally better than chance at p < 0.08, corrected for multiple comparisons within the respective time window. C. ERPs for visible/invisible flashes presented in the right or left locations. Grand averages were computed by averaging all trials for each condition first within each participant, then across participants. D. Topographies of ERP difference between flash right and flash left conditions, for visible (left) and invisible (right) trials. ERP topographies were averaged across 200–300 ms time window (i.e. the window that includes the maximal decoding accuracy, see figure 2B). In all figures: ***p < 0.001, **p < 0.01, *p < 0.05, ^[^*^]^p < 0.08, n.s. p > 0.08.

**Table 1.**
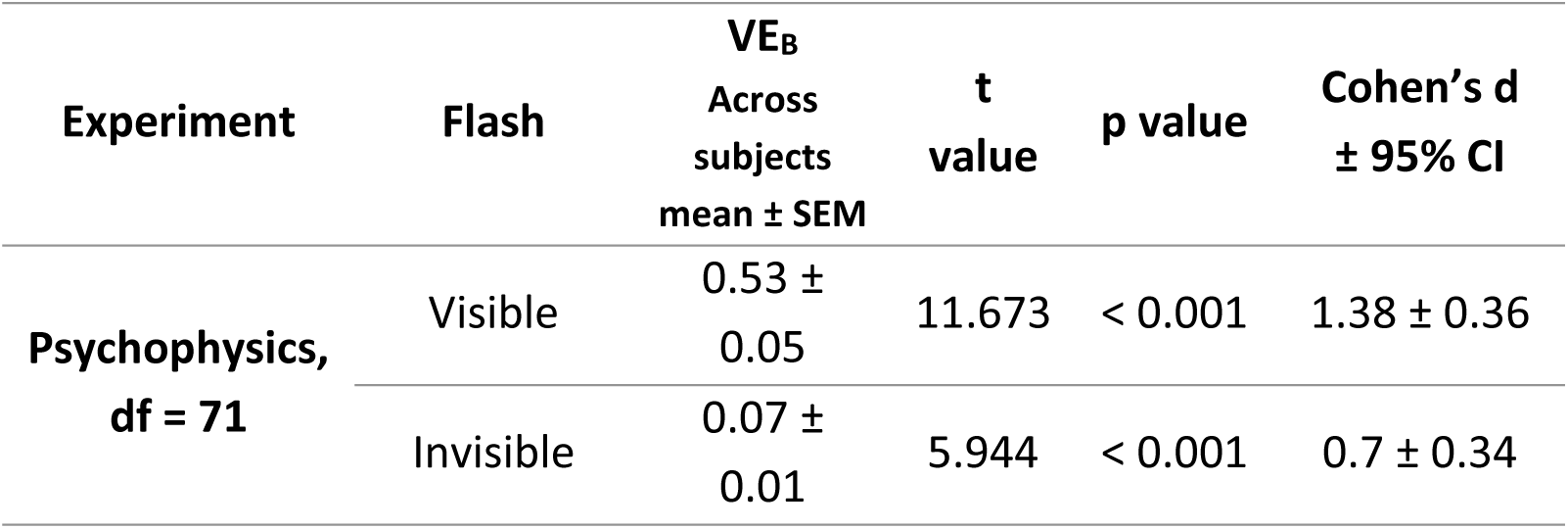
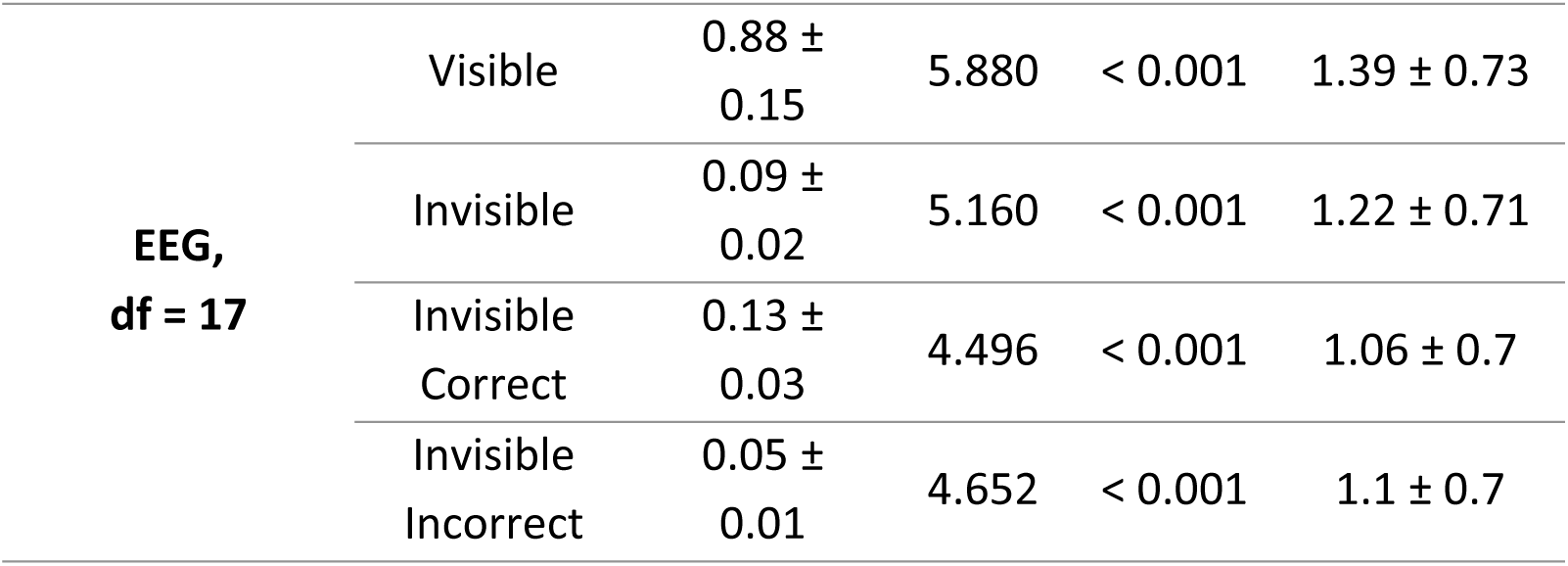
Behavioural results. Behavioural ventriloquist effect separately for trials on which the flash was i. visible or invisible and ii. correctly or incorrectly localized. t- and p-values refer to right tailed one sample t-tests. VE_B_ = behavioural ventriloquist effect.

| Experiment | Flash | VE<sub>B</sub> | t<br>value | p value | Cohen's d<br>± 95% CI |
| --- | --- | --- | --- | --- | --- |
|  |  | Across<br>subjects<br>mean ± SEM |  |  |  |
| Psychophysics,<br>df = 71 | Visible | 0.53 ±<br>0.05 | 11.673 | < 0.001 | 1.38 ± 0.36 |
|  | Invisible | 0.07 ±<br>0.01 | 5.944 | < 0.001 | 0.7 ± 0.34 |
| EEG,<br>df = 17 | Visible | 0.88 ±<br>0.15 | 5.880 | < 0.001 | 1.39 ± 0.73 |
|  | Invisible | 0.09 ±<br>0.02 | 5.160 | < 0.001 | 1.22 ± 0.71 |
|  | Invisible<br>Correct | 0.13 ±<br>0.03 | 4.496 | < 0.001 | 1.06 ± 0.7 |
|  | Invisible<br>Incorrect | 0.05 ±<br>0.01 | 4.652 | < 0.001 | 1.1 ± 0.7 |

Importantly, we observed a significant ventriloquist illusion for invisible trials even when subjects located the flash incorrectly (see Table 1, see Figure 2A). In other words, when observers mislocated an ‘invisible flash’, their perceived sound localization was on average biased towards the true (rather than reported) location of the flashes. In summary, the location not only of visible but also invisible and mislocalized flashes attracts observers’ perceived sound location and can induce a ventriloquist illusion.

Please note that this sub-analysis was only possible for the EEG study, in which we acquired an extensive data set over four days for each observer. For the single day psychophysics study, there were not sufficient trials on which observers mislocated the invisible flash. Likewise, it was not possible to assess the ventriloquist effect for visible flashes that were mislocated, because observers reliably rated their visibility and hence located almost all visible flashes correctly.

#### Control analysis: Effect of task order on the ventriloquist effect

In both the psychophysics and the EEG study we counterbalanced the task order (either: sound location – visibility – flash location or: flash location – visibility – sound location) across participants to account for the fact that the order of the questions may influence observer’s performance accuracy and response choices on the different tasks. For instance, as a result of memory noise, the order of questions may influence observers’ flash localization accuracy and the size of the ventriloquist effect. This may in turn affect the number of participants that can be included in the analysis. In the psychophysics experiment the number of included participants for each of the two question order was identical. In the EEG experiment, the proportion of included participants with either question order did not differ from the psychophysics experiment (chi^2^ = 0.178, p = 0.6732).

Moreover, one may argue that the ventriloquist effect for invisible flashes arises mainly on trials, when observers report the sound location before the flash location because memory noise impedes their visibility rating and flash localization performance. Contrary to this conjecture, two-tailed independent sample t-tests did not indicate a significant influence of question order on flash localization accuracy for visible (t(16) = 0.121, p = 0.905; Bayesian statistics: BF_01_ = 2.407) or invisible trials (t(16) = -0.056, p=0.956, BF_01_ = 2.417). Likewise, question order did not significantly influence the ventriloquist effect for visible (t(16) = 1.659, p = 0.117, BF_01_ = 0.978) and invisible trials(t(16) = 0.241, p = 0.813, BF_01_ = 2.371), whether the participants correctly (t(16) = 0.018, p = 0.986, BF_01_ = 2.419) or incorrectly (t(16) = 0.464, p = 0.649, BF_01_ = 2.246 ) localized the flash. Collectively, these control analyses confirm that spatial ventriloquism can arise for invisible flashes that are mislocated by observers irrespective of question order.

#### Relationship between the behavioural ventriloquist effect for visible and invisible flashes

The observation that both visible and invisible flashes can induce spatial ventriloquism raises the question whether these crossmodal biases are mediated via shared or different neural circuities. If visible and invisible flashes bias observers’ perceived sound location via partially shared mechanisms, we may expect that the magnitude of ventriloquist effects for visible and invisible flashes are correlated over participants. Indeed, we observed a significant correlation between observers’ ventriloquist effect for visible and invisible flashes (psychophysics experiment: Pearson’s R = 0.404, p < 0.001, EEG experiment: Pearson’s R = 0.73, p < 0.001). Participants that frequently experienced ventriloquist illusions for visible flashes also reported ventriloquist illusions more often for invisible flashes. This correlation provides initial suggestive behavioural evidence that the impact of visible and invisible flashes on sound processing may rely on partially shared neural mechanisms.

In summary, our behavioural results demonstrate that visible and invisible flashes elicit a robust ventriloquist effect possibly via partially shared mechanisms. Crucially, while this ventriloquist effect was unsurprisingly greater for visible than invisible flashes, we also observed a robust ventriloquist effect even for invisible flashes that were mislocalized by observers.

### EEG results

Combining EEG and multivariate pattern decoding we investigated the neural mechanisms that mediate the influence of the flash’s location on observers’ perceived sound location. First, we investigated how neural activity encodes the location of visible and invisible flashes that are correctly or incorrectly localized. Second, we assessed how this visuospatial information influences the neural encoding of sound location depending on the flash’s visibility, observers’ flash localization accuracy and experience of a perceptual ventriloquist illusion. We performed all analyses i. pooled over the entire time window from 0 to 650 ms and ii. temporally resolved across post-stimulus time using 50 ms sliding time windows.

#### Decoding flash location depending on its visibility and observers’ localization accuracy (complete time window)

To decode left vs. right flash location we trained a linear support vector machine (SVM) based on EEG signals from a 0–650 ms time window of audiovisual trials separately for visible and invisible flashes. Because we counterbalanced the sound locations across left and right flash classes (i.e. the number of left, middle and right sound locations was equal for left and right flash classes), the discriminative activity patterns were not confounded by sound location, but selective for the visuospatial information.

Classification accuracy for left vs. right flash location was significantly better than chance both for visible and invisible trials (see Table 2 for detailed results). To formally assess the similarity of neural representations elicited by visible and invisible flashes we generalized the classifier that was trained on visible trials to invisible trials. Again, ‘left vs. right flash’ classification accuracy was significantly better than chance and even slightly greater than the accuracy obtained for SVMs trained on invisible trials – though this improvement was not statistically significant (two-tailed paired t-test: t(17) = - 1.164, p = 0.261; Bayesian statistics: BF_01_ = 2.290). These results confirm that invisible flashes elicit similar, but weaker and/or less reliable neural representations than visible flashes. Indeed, visual inspection of the ERP topographies indicates that invisible flashes elicit ERPs with timecourses and topographies that are similar but attenuated relative to those for visible flashes (Figure 2C, D).

**Table 2.** EEG multivariate pattern analysis results: Classification accuracy for right vs. left flash location based on linear SVM tested/trained on EEG patterns [0 650] ms of audiovisual trials as specified in the table. t- and p-values refer to right tailed one sample t-tests against chance decoding accuracy of 50%.

| Tested / Trained | Decoding | t(17) | p value |
| --- | --- | --- | --- |
| | Accuracy<br>Across subjects<br>mean ± SEM % | | |
| Visible / Visible | 65.9 ± 2.5% | 6.383 | < 0.001 |
| Invisible / Invisible | 53.11 ± 0.9% | 3.500 | < 0.001 |
| Invisible / Visible | 54.2 ± 1% | 4.073 | < 0.001 |
| Invisible Correct / Visible | 54.6 ± 1.1% | 4.292 | < 0.001 |
| Invisible Incorrect / Visible | 53.7 ± 1.1% | 3.359 | < 0.001 |

Next, we focused on the invisible trials and asked whether SVMs decoding accuracy is reduced for invisible trials that are not correctly localized by observers. Contrary, to this conjecture, SVMs trained on visible trials successfully decoded the location of invisible flashes, even when the flashes were mislocalized by observers (see Table 2 for complete results). Further, we did not observe a significant difference in classifier performance for invisible flashes that were correctly or incorrectly localized (two-tailed paired t-test: t(17) = 0.624, p = 0.541; Bayesian statistics: BF_01_ = 3.460).

#### The neural dynamics of flash location encoding depending on its visibility and observers’ localization accuracy (temporally resolved analysis)

We temporally resolved the neural encoding of flash location by training SVMs to classify left vs. right flashes (again counterbalanced for sound location) based on EEG activity patterns confined to 50 ms windows sliding across poststimulus time. We successfully (i.e. significantly better than chance) discriminated between left and right flashes on visible trials from about 160 ms poststimulus until the end of the 650 ms time window. Likewise, for invisible flashes we obtained better than chance classification accuracy from 200 ms onwards till the end of the time window when observers located the flash correctly (see Figure 2B for details). This was true for SVM classifiers trained on visible as well as invisible trials indicating that visible and invisible flashes were at least partly encoded via shared neural circuitries. The time course of decoding accuracy had a similar temporal profile for visible and invisible flashes, peaking at about 230-250 ms followed by a rapid decline in decoding accuracy. Yet, overall, the decoding accuracy for visible flashes was markedly higher than that for invisible flashes, even when observer located them correctly. The decoding accuracy and neural activity patterns for visible and invisible flashes were related directly to observers’ visibility ratings.

Moreover, between 330 and 530 ms post stimulus the traces of decoding accuracy diverged and were significantly different for correctly and incorrectly localized invisible flashes (see Figure 2B). The decoding accuracy for correctly located invisible flashes was significantly above chance throughout the entire time window, but rapidly decayed to chance accuracy after 330 ms for incorrectly located invisible flashes. These results show that visuospatial information encoded in EEG activity pattern is directly related to whether observers’ ability to locate an invisible flash correctly. Only invisible flashes that observers correctly located evoked EEG activity patterns that enabled sustained above chance decoding accuracy across the entire time window.

Collectively, these decoding results demonstrate that the location of visible and invisible flashes is initially encoded in similar EEG activity patterns though less reliably for invisible flashes. Critically, when the flash is invisible and incorrectly localized by observers, spatial information about the flash rapidly dissipates from the visual system at about 330 ms post stimulus as indicated by a significant decoding difference for correctly and incorrectly located invisible flashes.

#### The neural ventriloquist effect: crossmodal spatial biases for visible and invisible flashes (complete time window analysis)

To investigate how visible and invisible flashes influence the neural encoding of sound location we trained a support vector regression (SVR) model to learn the mapping from EEG activity patterns (0 to 650 ms) to external sound location (left, middle, right) based on unisensory auditory trials alone (i.e. the three no flash conditions). This SVR model is therefore predominantly sensitive to auditory spatial information. We then used this trained SVR model to predict the sound location from EEG activity patterns of audiovisual trials. If the location of a synchronous flash biases the neural encoding of sound location, we would expect that the decoded sound location on audiovisual trials is shifted towards the flash location.

To assess this flash-induced spatial bias we computed the so-called neural ventriloquist effect as the average decoded sound location for ‘visual right’ minus ‘visual left’ trials (i.e. exactly as we did in the behavioural analysis for reported sound locations). Figure 3B shows the sound location estimates decoded from audiovisual EEG patterns within the entire [0 650] ms time window separately for trials on which the flash was judged visible and invisible. Consistent with the behavioural results, the neural ventriloquist effect was significant for both visible and invisible trials, albeit ∼5 times smaller than the behavioural ventriloquist effect. As expected, the decoded sound location shifted towards the location of the concurrent flash.

**Figure 3.**
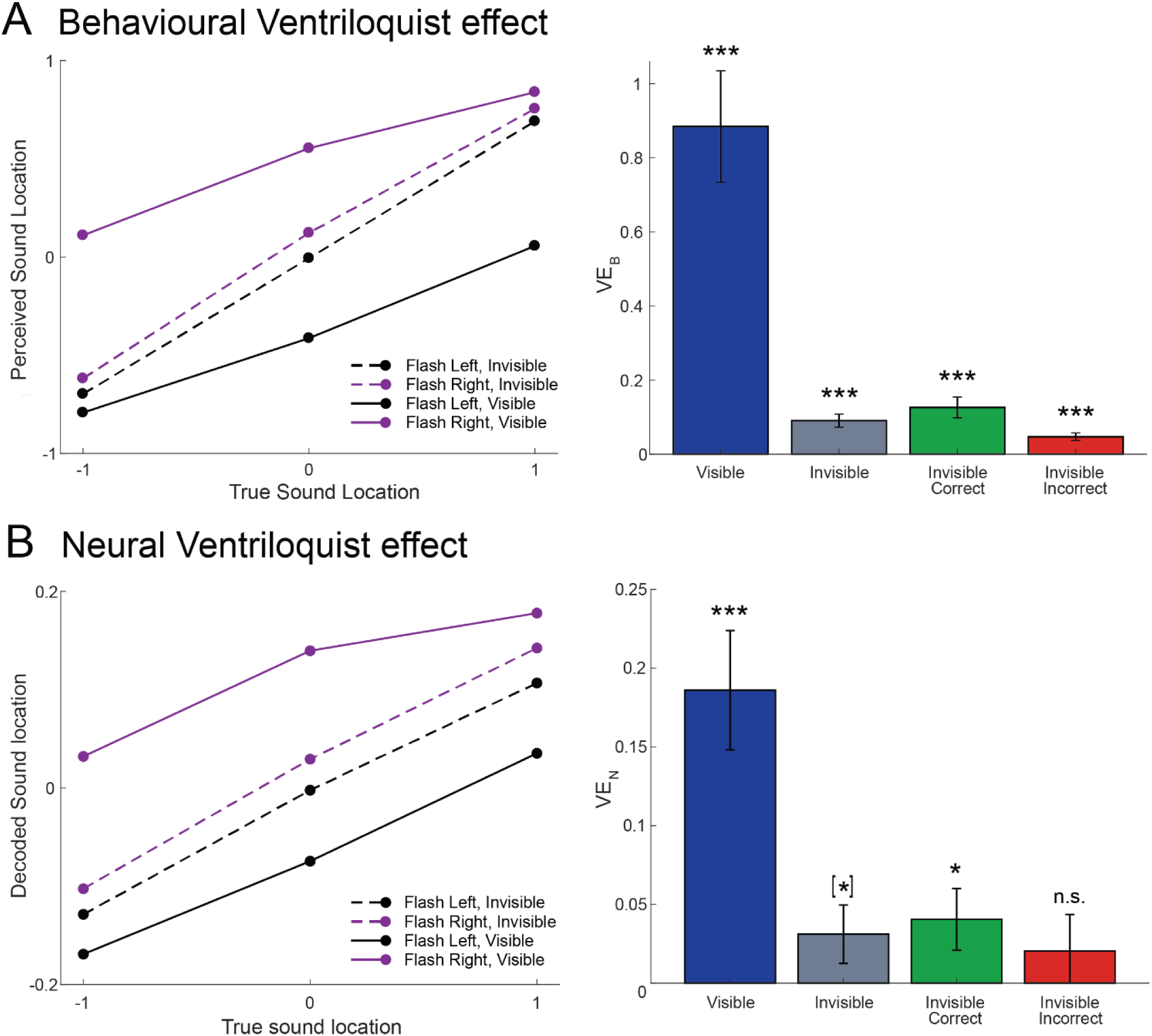
Behavioural and neural ventriloquist effect. A. Behavioural data: Reported sound locations (across subjects mean) for AV trials and behavioural ventriloquist effect (VE_B,_ across subjects mean ± SEM). B. EEG data: Sound locations (across subjects mean) decoded from EEG activity patterns across [0 650] ms time window poststimulus and neural ventriloquist effect (VE_N_, across subjects mean ± SEM).

Mimicking the behavioural results, the ventriloquist effect was attenuated for invisible relative to visible trials (two-tailed paired t-test t(17) = 5.191, p < 0.001 ; BF_01_=0.003, see Figure 3B). It was significantly greater than zero for invisible trials that were correctly localized (see Table 3).

**Table 3.** EEG results (complete time window analysis): Neural ventriloquist effect separately for trials on which the flash was visible/invisible and/or correctly/incorrectly located by observers. t- and p-values refer to two-tailed one sample t-tests. VE_N_ = neural ventriloquist effect.

| Flash | VE<sub>N</sub><br>Across<br>subjects<br>mean ± SEM | t(17) | p value | Cohen’s d<br>± 95% CI |
| --- | --- | --- | --- | --- |
| Visible | 0.19 ± 0.04 | 4.906 | < 0.001 | 1.16 ± 0.71 |
| Invisible | 0.03 ± 0.02 | 1.676 | 0.056 | 0.39 ± 0.66 |
| Invisible<br>Correct | 0.04 ± 0.02 | 2.067 | 0.027 | 0.49 ± 0.66 |
| Invisible<br>Incorrect | 0.02 ± 0.02 | 0.881 | 0.195 | 0.21 ± 0.65 |

#### The relationship between the behavioural and the neural ventriloquist effect for visible and invisible flashes (complete time window analysis)

Next, we assessed the relationship between the neural and behavioural ventriloquist effect. For this, we computed the neural ventriloquist effect for invisible and visible flashes separately for trials on which observers did or did not experience a perceptual ventriloquist illusion (n.b. this analysis can be performed in an unbiased fashion only for sound centre trials, see Methods section). We observed a highly significant ventriloquist effect when observers experienced a ventriloquist illusion both for visible and invisible flashes (see Figure 4A, Table 4, two-tailed one-sample t-tests against zero). By contrast, the neural ventriloquist was not significantly greater zero when observers did not experience a perceptual ventriloquist illusion - though we acknowledge a non-significant trend for visible flashes without a behavioural ventriloquist effect.

**Figure 4.**
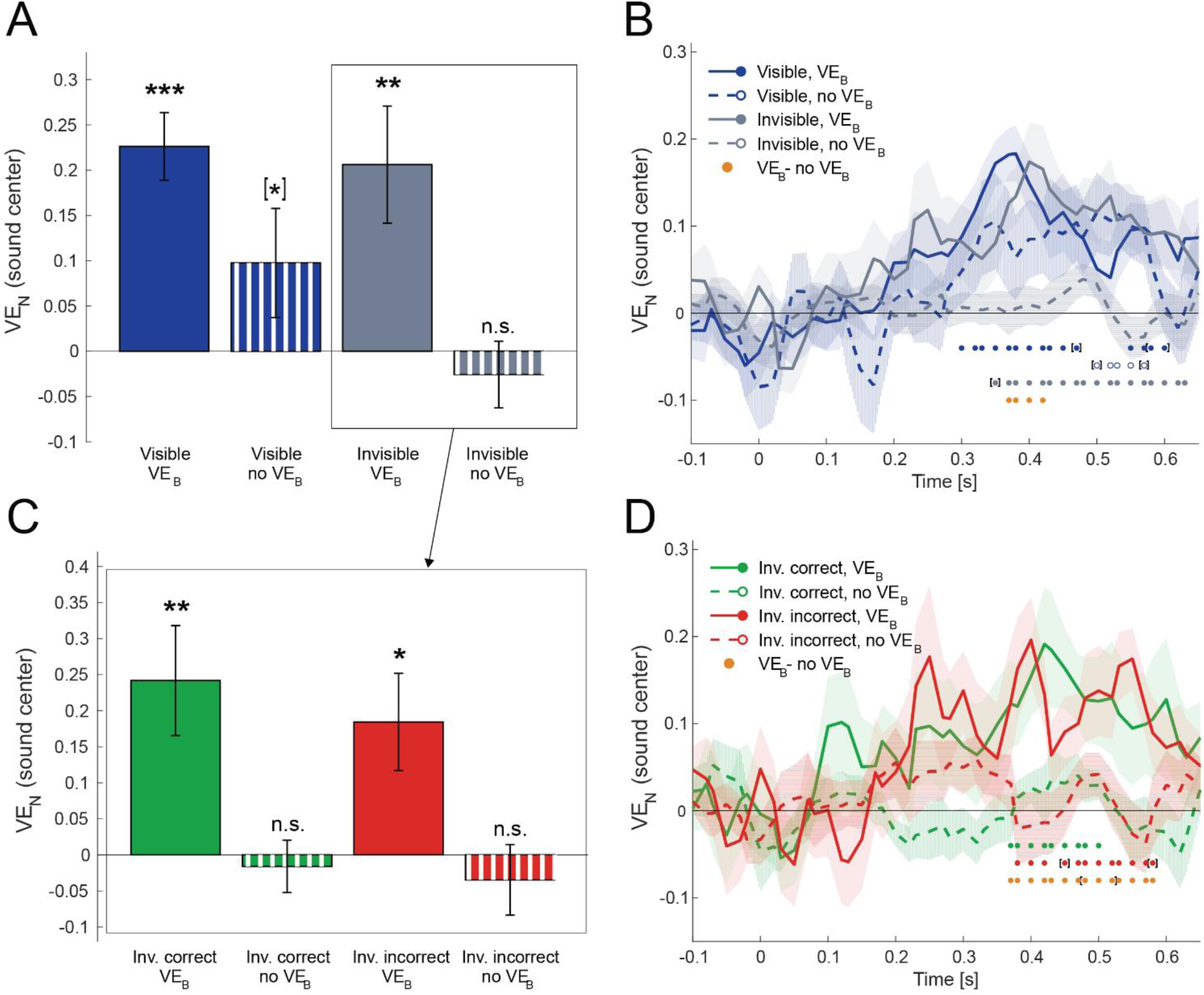
Neural ventriloquist illusion for sound centre location, for trials with or without reported ventriloquist illusion (VE_B_). A,C Bar plots showing VE_N_ (mean ± SEM) for classifier predictions based on the entire time window [0-650] ms for visible and invisible flashes (A) and correctly/incorrectly located invisible flashes (C). B,D Time course of neural ventriloquism (VE_N_) for visible and invisible flashes (B) and correctly/incorrectly located invisible flashes (D). Dots indicate VE_N_ significantly different than zero at p < 0.05 corrected for multiple comparisons within the [0 650] ms time window. Temporal smoothing (average over 2 neighbouring data points) was applied to plots in B&D for visualization purposes (statistical analyses were performed for non-smoothed data).

**Table 4.** EEG results (complete time window analysis): Neural ventriloquist effect selectively for sound centre trials reported separately for trials on which the flash was i. visible/invisible, ii. correctly/incorrectly located by observers and iii. a behavioural ventriloquist was present/absent (VE_B_= yes/no). t- and p-values refer to two-tailed one sample t-tests. VE_N_ = neural ventriloquist effect. VE_B_ = behavioural ventriloquist effect.

| Flash | VE<sub>B</sub> | VE<sub>N</sub><br>Across<br>subjects<br>mean ± SEM | t(17) | p value | Cohen's d<br>± 95% CI |
| --- | --- | --- | --- | --- | --- |
| Visible | yes | 0.23 ± 0.04 | 6.044 | **< 0.001** | 1.42 ± 0.73 |
| | no | 0.1 ± 0.06 | 1.616 | 0.062 | 0.38 ± 0.66 |
| Invisible | yes | 0.21 ± 0.06 | 3.184 | **0.003** | 0.75 ± 0.68 |
| | no | -0.03 ± 0.04 | -0.700 | 0.753 | -0.16 ± 0.65 |
| Invisible<br>Correct | yes | 0.24 ± 0.08 | 3.172 | **0.006** | 0.75 ± 0.68 |
| | no | 0.02 ± 0.04 | -0.442 | 0.664 | -0.10 ± 0.65 |
| Invisible<br>Incorrect | yes | 0.18 ± 0.07 | 2.737 | **0.014** | 0.64 ± 0.67 |
| | no | -0.03 ± 0.05 | -0.709 | 0.488 | -0.17 ± 0.65 |

Consistent with these one-sample t-test results, a 2(visibility: visible vs. invisible) x 2(behavioural ventriloquist effect: present vs. absent) repeated measures ANOVA revealed a significant effect of behavioural ventriloquism on neural VE (F(1,17) = 20.146, p = < .001), but not of visibility (F(1,17) = 2.312, p = 0.147) and no interaction (F(1,17) = 0.973, p = 0.338).

Next, we focused on the neural ventriloquist effect for invisible flashes. We assessed whether the neural ventriloquist effect for invisible flashes differed for invisible flashes that observers localized correctly or incorrectly (see Figure 4C, Table 4). Again, this analysis highlighted the tight relationship between the neural and the behavioural ventriloquist effects: we observed a significant neural ventriloquist for both correctly and incorrectly localized invisible flashes only, when observers experienced a ventriloquist illusion. By contrast, no significant neural ventriloquist effect was observed, when the illusion did not occur. A 2 (flash localization accuracy: correct vs. incorrect) x 2(behavioural ventriloquist effect: present vs. absent) repeated measures ANOVA revealed an effect of behavioural ventriloquism (F(1,17) = 14.063, p = 0.002), but not of flash localization accuracy (F(1,17) = 0.817, p = 0.379) and no interaction (F(1,17) = 0.203, p = 0.658).

Collectively, these results demonstrate that the neural ventriloquist effect is closely related to the behavioural ventriloquist effect. When observers experience a ventriloquist illusion, the magnitude of the neural ventriloquist effect (i.e. the impact of flash location on decoded sound location) is comparable for all flashes irrespective of their visibility or localizability. This contrasts with the results we obtained for the flash decoding, showing a difference in decoding accuracy for visible and invisible flashes, even when observers located the flash correctly. In other words, flash visibility had a profound impact on flash decoding accuracy even if observers’ behavioural flash location responses were matched. By contrast, flash visibility did not have a significant effect on the neural ventriloquist effect.

#### Influence of question order on decoding accuracy of flash location and neural ventriloquism

Consistent with our behavioural results, two-tailed independent samples t-test did not reveal a significant effect of question order on decoding accuracy of flash location (visible: p = 0.644, t(16) = 0.471, BF_01_ = 2.241, invisible: p = 0.904, t(16) = 0.123 , BF_01_ = 2.407, invisible trained visible: p = 0.578, t(16) = - 0.568, BF_01_ = 2.164, invisible correct trained visible: p = 0.752, t(16) = -0.321, BF_01_ = 2.334, invisible incorrect trained visible: p = 0.684, t(16) = 0.414, BF_01_ = 2.280). Likewise, we did not observe a significant effect of question order on the neural ventriloquist illusion (visible: p = 1, t(16) = < 0.001, BF_01_ = 2.420, invisible: p = 0.752, t(16) = 0.321, BF_01_ = 2.334, invisible correct: p = 0.583, t(16) = 0.560, BF_01_ = 2.171, p = 0.869, t(16) = 0.168, BF_01_ = 2.396). Collectively, these null-results suggest that the question order cannot explain the flash decoding accuracies or the neural ventriloquist effect. Most importantly, the neural ventriloquist effect for invisible and mislocalized flashes was not significantly greater when observers first indicated their perceived sound location followed by visibility and flash localization that may have been impeded by additional memory noise.

#### Relationship between the neural ventriloquist effect for visible and invisible flashes

Consistent with our behavioural results, we observed a significant correlation between observers’ ventriloquist effect for visible and invisible flashes (Pearson’s R = 0.63, p = 0.005). Participants whose spatial encoding of sounds was more biased by visible flash location also showed greater bias of neural spatial encoding of the sound for invisible flashes. This correlation provides initial suggestive neural evidence that the impact of visible and invisible flashes on sound processing may rely on partially shared mechanisms.

#### The temporal dynamics of the neural ventriloquist effect for visible and invisible flashes (temporally resolved analysis)

We temporally resolved the neural ventriloquist effect by training support vector regression models on the unisensory auditory EEG activity patterns within 50 ms windows sliding across poststimulus time. Temporally resolved decoding revealed that the neural ventriloquist effect emerged around 300 ms for visible and 360 ms for invisible flashes when observers experienced the perceptual ventriloquist illusion (Figure 4B), the neural ventriloquist peaked at a similar time about 380-400 ms. For visible flashes, we also observed a significant neural ventriloquist effect, peaking later at about 500 ms, when observers did not experience a perceptual ventriloquist illusion. By contrast, no significant neural ventriloquist effects was observed for invisible flashes in the absence of a ventriloquist illusion. As a result of those different time courses, we observed a significant main effect of the behavioural ventriloquist illusion starting at about 360 ms post stimulus and ending 410 ms post stimulus, with a peak at 380 ms. We observed no main effect of visibility and no interaction of visibility and the behavioural ventriloquist effect (see Table 5).

**Table 5.** EEG results (time course analysis): Significant results (p < 0.05, corrected at the cluster level). Abbreviations: VE_B_, behavioural ventriloquist effect; Vis, Visible trials; Invis, Invisible trials; FLA, flash localization accuracy; I_C, Invisible correct trials; I_I, Invisible incorrect trials. Consecutive samples with significant results are grouped in the same time window and matched with a range of p-values corresponding to it.

| Effect | Neural |
| --- | --- |
| **Visibility** | n.s. |
| **VE<sub>B</sub> in Vis/Invis** | 366 - 416ms: 0.0093<p<= 0.0370 |
| **Visibility \* VE<sub>B</sub> in Vis/Invis** | n.s. |
| **FLA** | n.s. |
| **VE<sub>B</sub> in I_C and I_I** | 366-466ms: 0.009<p<0.0449<br>533-583ms : 0.0026<p<0.0406 |
| **FLA \* VE<sub>B</sub> in I_C and I_I** | n.s |

Focusing on invisible flashes alone, we next compared the neural ventriloquist effect for correctly and incorrectly localized invisible flashes (Figure 4D). Dovetailing nicely with our behavioural and complete time window results, we observed a significant neural ventriloquist effect irrespective of observers’ flash localization accuracy when observers experienced a ventriloquist illusion – but not otherwise. Likewise, the build-up of the neural ventriloquist effect was indistinguishable for correctly and incorrectly located invisible flashes, in both cases peaking at about 400 ms. Conversely, the neural ventriloquist effect differed significantly for trials with and without behavioural ventriloquist effect also from about 360–580 ms, peaking at 400 ms. We observed no main effect of flash localization accuracy and no interaction of the latter with the presence of a behavioural ventriloquist illusion.

In summary, temporally resolved decoding confirmed that the flash location biased spatial encoding of sounds irrespective of flash visibility or observers’ localization accuracy on trials when observers experienced the ventriloquist illusion. For trials without a behavioural ventriloquist effect, only visible but not invisible flashes induced a neural ventriloquist effect.

## Discussion

This psychophysics-EEG study combined dynamic continuous flash suppression (Maruya et al., 2008; Tsuchiya & Koch, 2005) and spatial ventriloquism to investigate how flashes that evade our subjective visual awareness influence the neural encoding and perception of sound location.

Consistent with previous behavioural results (Delong et al., 2018), observers perceived the sound shifted (i.e. biased) towards the true location of both visible and invisible flashes. While spatial ventriloquism was more frequently observed for visible flashes, a robust ventriloquist effect was also observed for invisible flashes. Critically and going beyond previous findings, we also observed a significant behavioural ventriloquist effect for invisible flashes that were mislocated by observers. In other words, observers’ perceived sound location was biased towards the true flash location, even when this visual input did not enable accurate spatial classification in vision (Figure 3). This ventriloquist illusion for invisible flashes contrasts with the McGurk illusion that is known to falter when visual inputs are obliterated from consciousness (Ching et al., 2019; Palmer & Ramsey, 2012). The discrepancy may be explained by the fact that the ventriloquist illusion relies on spatiotemporal binding, while the McGurk illusion requires integration of complex audiovisual features such as visemes (i.e. articulatory facial movements) and phonemes that are known to be more strongly impaired when awareness of sensory inputs is experimentally suppressed (Heyman & Moors, 2014; Moors et al., 2016). Evidence from research on processing of object-scene relations corroborates the absence of integration of complex stimuli without awareness (Biderman & Mudrik, 2018). Consistent with this conjecture, a recent study has shown that spatial but not semantic correspondence cues influence audiovisual binding in the absence of subjective awareness (Delong & Noppeney, 2021).

At the neural level, we first assessed how the brain encodes the location of a flash depending on observers’ visibility rating and flash localization accuracy. As shown in Figure 2, the location of a flash was decoded successfully from early (150-250 ms) neural activity for visible and invisible flashes - albeit a greater decoding accuracy was obtained for visible flashes. This is not surprising, because invisible flashes elicited ERPs with topographies that mimicked the topographies for visible flashes in an attenuated or less reliable fashion (Figure 2C & D). Crucially, when an invisible flash was mislocated by observers, spatial information rapidly dissipated from the visual system and was no longer decodable from EEG activity after 330 ms.

The extinction of subjectively unaware visuospatial information from EEG activity seemingly contradicts a recent study that was able to decode the location of invisible stimuli successfully from MEG activity until 800 ms, even when observers were not able to locate the stimuli accurately (Salti et al., 2015). However, contrary to this previous work, in our study the left/right visuospatial information that our EEG decoding focused on was task-irrelevant, because observers were engaged in an orthogonal up/down spatial classification task. Consistent with this explanation, King et al. (2016) have shown that only task-relevant information (e.g. orientation of a Gabor patch) was maintained until 1400 ms, while task-irrelevant information (e.g. spatial frequency) was no longer decodable from MEG activity after 230 ms poststimulus (King et al., 2016).

Collectively, the behavioural and neural results discussed so far pose the critical question of how ‘unaware’ visuospatial (left vs. right) information, that evaporates from the visual system at about 300 ms, can bias the neural encoding and perception of sounds.

To address this question we trained a support vector regression model to learn the mapping from unisensory auditory EEG activity to external sound location. This training scheme ensured that the decoding was selectively sensitized to the auditory processing system. We then used this mapping to decode the sound location from EEG activity of audiovisual trials. Consistent with our behavioural results the location of a flash biased the sound location that was decoded from EEG activity patterns. This neural ventriloquist effect closely paralleled the behavioural ventriloquist effect: it was significant for visible, invisible and even mislocalized invisible flashes on trials with a behavioural ventriloquist effect. When observers experienced a ventriloquist illusion, the neural ventriloquist effect emerged with a similar timecourse irrespective of observers’ visibility ratings: it started to rise at about 150 ms and culminated around 380 ms poststimulus for visible and invisible flashes, even when observers mislocalized them (Figure 4, see also (Bonath et al., 2007)). By contrast, in the absence of a behavioural ventriloquist effect only visible, but not invisible flashes, biased the neural encoding of sounds at a slightly later time. The emergence of a neural ventriloquist effect in the absence of a behavioural ventriloquist effect suggests that it truly reflects visual induced biases of sound encoding rather than later response preparation processes. In fact, if the neural ventriloquist effect reflected response preparation processes, it should not peak at 300-400 ms for invisible flashes, but constantly rise until the presentation of the response cue at 650 ms. Also response preparation processes should depend on question order. By contrast, the neural and behavioural ventriloquist did not significantly differ for trials on which observers initially reported the sound vs. the flash location.

Our results thus suggest that unconscious visuospatial information that dissipates from the visual system at about 300 ms continues to have a sustained impact on spatial coding in the auditory system from 150 ms onwards, thereby ultimately inducing a perceptual ventriloquist illusion. The common timecourses suggest that these audiovisual interactions rely on partially shared neural mechanisms and circuities for visible and invisible flashes – thereby also explaining their tight correlation in observers’ tendency to experience a ventriloquist illusion.

Our current findings dovetail nicely with recent EEG (Aller & Noppeney, 2019) and fMRI (Rohe & Noppeney, 2015a, 2016, 2018) research that spatiotemporally resolved integration of supraliminal audiovisual spatial signals progresses across the cortical hierarchy. Consistent with the current results, this previous research showed that audiovisual signals are integrated into spatial representations starting at about 150 ms in sensory cortices and culminating at about 300-500 ms in higher order areas such as the planum temporale and parietal cortices (see also (Bonath et al., 2007)). In the light of this prior work and our flash location decoding results, invisible flashes may influence spatial encoding of sounds via early interactions that rely on direct connectivity between sensory cortices (Foxe & Schroeder, 2005; Ghazanfar & Schroeder, 2006) ultimately leading to a neural and a perceptual ventriloquist effects that are indistinguishable from those observed for visible flashes. We propose that the spatial information present in the visual system can thus ascend the cortical hierarchy via the auditory cortex ((Bieler et al., 2018; Komura et al., 2005; Vittek et al., 2023);see Figure 5 showing proposed audio-visual integration model). A final common pathway for ventriloquism in the absence and presence of subjective awareness is also supported by an across-participants correlation between the visible and invisible ventriloquist effects. Participants that frequently experienced the ventriloquist illusion for visible flashes also reported ventriloquist illusions more often for invisible flashes, again suggesting at least partially shared mechanisms.

**Figure 5.**
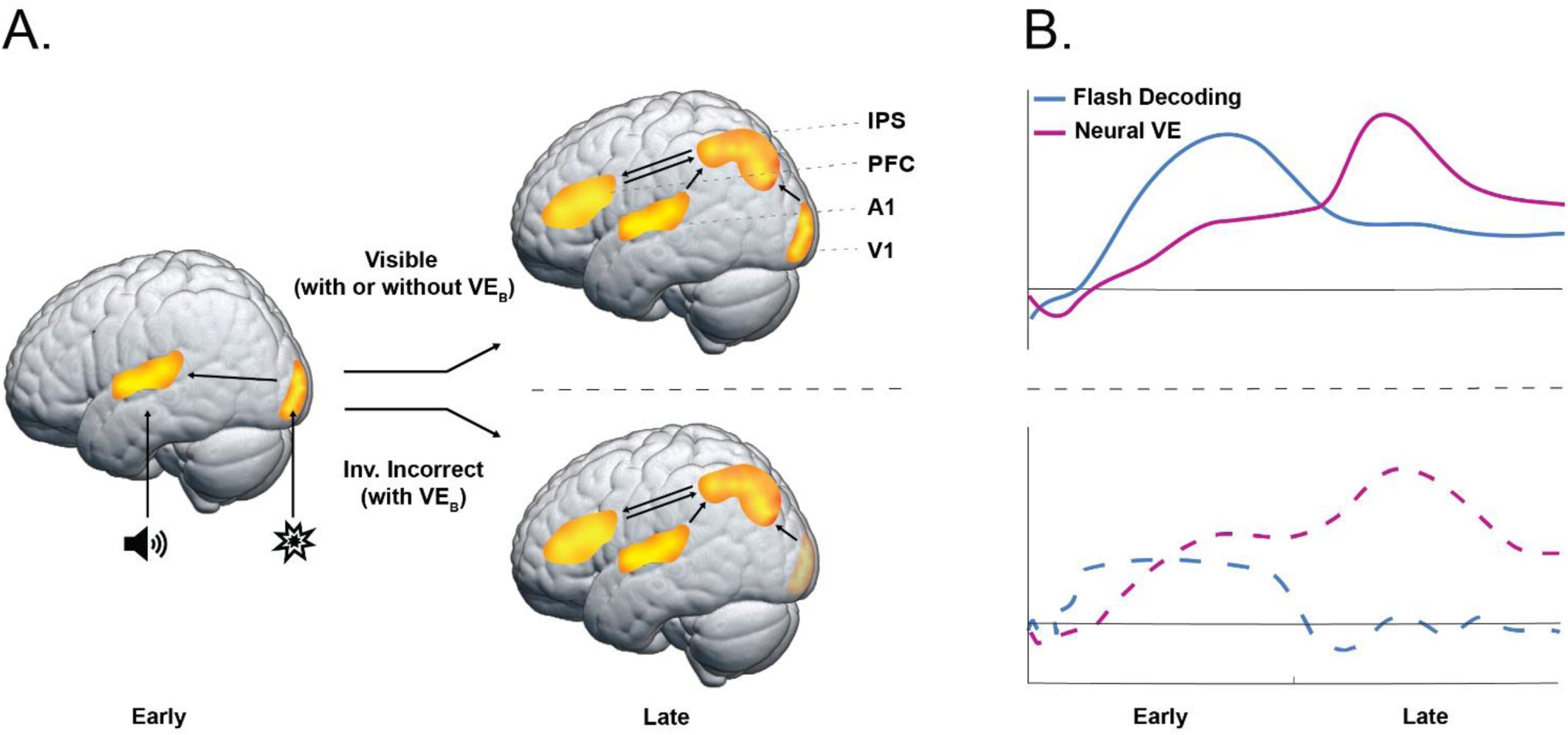
Mechanisms of audiovisual interactions in ventriloquism. (A). Mechanisms of how invisible flashes can influence sound perception. On trials on which the flash is visible, visual spatial information is available in early (left) and late (top right) processing stages to influence neural spatial representations of sound. On trials on which the flash is invisible and incorrectly localized by observers, visual spatial information dissipates in the visual system (bottom right) at about 330ms, but can nonetheless influence auditory spatial processing via earlier interactions (left), so that the neurally encoded and reported sound location is shifted towards the true flash location (i.e. neural and behavioural ventriloquist effect). (B). Time course of decoded flash location accuracy (in blue) and of the neural ventriloquist effect (in purple) for visible (top) and for incorrectly localized invisible trials (bottom). For visible trials both the flash decoding accuracy and neural VE remain during the late processing stages. For incorrectly localized invisible trials, flash decoding accuracy decreases to chance level after early processing stages while the neural VE remains. Spatial locations: L = left, C = centre, R = right. VE = ventriloquist effect

Our results challenge current leading theories that consider consciousness as an ‘all or none’ phenomenon which arises when signals enter the Neuronal Global Workspace via non-linear process of ignition. We show that visuospatial information that dissipates from the visual system and evades visual awareness impacts neural encoding and conscious auditory perception. In other words, visuospatial information that is not accessible to visual awareness is partly preserved in the spatial representations of consciously perceived sounds as a result of audiovisual integration.

The dissociation between the fate of invisible flashes in the visual system and their sustained impact on spatial encoding in the auditory system can be explained by models that treat perception as probabilistic inference based on noisy sensory inputs (Körding et al., 2007; Ma, 2012). Importantly, internal noise can influence information transmission partly independently along different sensory processing hierarchies (Faisal et al., 2008; Van Vugt et al., 2018). Indeed, a recent elegant neurophysiological study in awake monkeys has demonstrated that noise may prevent the propagation of information along the visual processing hierarchy at various stages, thereby obliterating visual inputs from perceptual awareness (Van Vugt et al., 2018). Multisensory processing provides the additional possibility that noise interferes with representations selectively in the auditory or visual processing streams. As a result, visual inputs can bias the spatial encoding of sounds via early (<300 ms) multisensory interactions, even if additional noise along the visual processing hierarchy prevents this visuospatial information from entering observers’ visual awareness. In short, internal noise or stochastic effects in the neural system turn consciousness into a heterogeneous phenomenon: information that eludes observers’ awareness in one sensory modality can impact neural encoding and ‘conscious’ perception in other sensory systems - posing new questions about the essence of multisensory awareness (Deroy et al., 2014, 2016). Interestingly, the decoding accuracy differed markedly for visible and invisible flashes even if the latter were correctly localized by observers presumably because of stochastic effects in decision making. By contrast, the neural ventriloquist effect arose with nearly identical time courses irrespective of flash visibility and localizability when observers experienced the perceptual ventriloquist illusion. These results suggest that stochastic effects determine whether visuospatial information interacts with auditory processing during early processing stages; yet, by being integrated with the strong supraliminal auditory input this visual information is less vulnerable to internal noise and preserved along auditory processing stream.

In conclusion, our findings unravel the neural mechanisms by which a flash that evades our awareness alters where we perceive sounds. ‘Unconscious’ visuospatial information that dissipates from the visual system within 300 ms poststimulus can continue to have a sustained impact on neural spatial encoding in the auditory system. These results demonstrate that consciousness is not a generic prerequisite for information integration, not even in situations in which signals stem from different sensory channels. Our results pose challenges for leading theories of consciousness as a unified ‘all-or-none’ phenomenon.

## Methods

### Participants

After giving informed consent, 103 (78 females, 8 left-handed, mean age: 21.5 years, standard deviation: 4.9, range: 18-41; 41 participants were included in (Delong et al., 2018)) healthy young adults took part in the psychophysics experiment; 72 of those participants were included in the analysis (see inclusion criterion section). Eighteen of those subjects participated (13 females, 2 left-handed, mean age = 21.2 years, standard deviation = 4.2, range: 18-31) in the subsequent EEG experiment. The study was performed in accordance with the principles outlined in the Declaration of Helsinki and was approved by the local ethics review board of the University of Birmingham.

To determine the sample size for the EEG experiment we used the effect size (Cohen’s d ≈ 0.7) for the ventriloquist effect for invisible trials based on the psychophysics studies described in Delong et al. 2018. For statistical power of 0.9 we obtained n = 18. We continued acquiring subjects for the psychophysics experiment until the number of the EEG data sets was equal to the required sample size (i.e. excluded subjects were replaced; see Inclusion criteria section).

### Inclusion criteria for psychophysics and EEG

The study included a one day psychophysics experiment and four day EEG experiment (for a subset of 18 subjects).

For the psychophysics experiment, we limited the analysis to 72 subjects based on two exclusion criteria: First, we included only subjects that provided reliable visibility judgments as indicated by ‘better than chance localization accuracy’ (based on binomial test) for visible flashes (i.e. exclusion of 22 subjects). Second, to ensure reliable parameter estimation, we included only those participants who had at least 10 trials in each of the 2 (flash location: left vs. right) x 3 (sound location: left, middle, right) conditions, for both visible and invisible categories respectively (i.e. exclusion of 9 subjects). These exclusion criteria ensured that the computation of the ventriloquist effect was based on at least 60 trials.

For the EEG experiment we invited subjects back from the initial psychophysics experiment, if their ventriloquist effect for invisible trials was equal to or greater than 0.05, and their flash localization accuracy for invisible trials was not higher than 56%. 31 out of 72 subjects were invited to participate in the EEG experiment, but only 18 completed all four EEG sessions. We included all participants that completed all four EEG sessions in the analysis.

### Stimuli and apparatus

Participants sat in a dimly lit room in front of a computer monitor at a viewing distance of 95 cm. They viewed one half of the monitor with each eye using a custom-built mirror stereoscope. Visual stimuli were composed of targets and masks that were presented on a grey, uniform background with a mean luminance of 15.6 cd/m^2^. On the ‘flash present’ trials, one eye viewed the target stimulus (i.e. the flash), which was a grey disc (Ø 0.3°) presented for 50ms in the upper left, lower left, upper right or lower right quadrant, i.e. at ± 3° visual angle along the azimuth and ± 1.2° elevation from a grey central fixation dot. The elevation of ± 1.2° was selected to enable effective multisensory interactions between flash and sound irrespective of flash elevation. The luminance of the flash was adjusted individually via adaptive staircases to obtain 60% invisible trials. To suppress the flash’s perceptual visibility, we used a dynamic version of continuous flash suppression (dCFS) in which four dynamic Mondrians (Ø 2.08°, mean luminance: 48 cd/m^2^) rather than rectangles (Tsuchiya & Koch, 2005) were shown to the other eye (Aller et al., 2015; Maruya et al., 2008). Each Mondrian consisted of sinusoidal square gratings (d = 0.6°) which changed their colour and position randomly at a frequency of 20Hz. Each grating’s texture was shifted every 16.6ms (i.e. each frame of the monitor with 60Hz refresh rate) to generate apparent motion. Visual stimuli were presented at four possible locations that were equidistant from a central fixation spot. They were framed by a grey aperture (thickness: 0.15°, luminance: 110 cd/m^2^) of 8.97° x 14.15° in diameter to aid binocular fusion. Mask and target screen allocation (right, left eye) alternated between eyes across trials, to enhance suppression.

Auditory stimuli were 50 ms bursts of white noise. They were presented via six external speakers, placed above and below the monitor at 64 dB sound pressure level. Upper and lower speakers were aligned vertically and located centrally, 3° to the left and 3° to the right of the monitor’s centre (i.e. aligned with the flash location along the azimuth).

Psychophysics stimuli were generated and presented using the Psychtoolbox version 3.0.11 (Brainard, 1997) on MATLAB R2014a (Mathworks, Natick, Massachusetts) with a PC running Windows XP. Staircase procedures were implemented using Palamedes toolbox (Prins & Kingdom, 2009).

Visual stimuli were presented dichoptically using a gamma-corrected 30” LCD monitor with a resolution of 2560 x 1600 pixels at a frame rate of 60Hz (NVIDIA Quadro 600 graphics card). Auditory stimuli were digitized at a sampling rate of 44.8 kHz via an M-Audio Delta 1010LT sound card. Exact audiovisual onset timing was confirmed by recording visual and auditory signals concurrently with a photodiode and a microphone.

### Experimental Design

In a spatial ventriloquist paradigm, participants were presented with an auditory burst of white noise emanating from one of three potential locations: left, centre or right. In synchrony with the sound, one eye was presented with i. no flash or a brief flash in participants’ ii. left or iii. right hemifield under dynamic continuous flash suppression to the other eye (Maruya et al., 2008). Hence, the 3 x 3 factorial design manipulated i. ‘flash ‘ (3 levels: left flash, right flash, no flash) and ii. ‘sound location’ (3 levels: left sound, central sound and right sound) (Figure 1A). In order to enable a flash localization task that is orthogonal to the sound localization, the flash could be presented either in the upper or lower hemifield (i.e. ± 1.2° elevation from a grey central fixation dot). Hence, the flash was presented in the upper left quadrant, lower left quadrant, upper right quadrant or lower right quadrant (n.b. visual localization is highly precise close to the fixation point and has been shown to be equivalent for spatial discrimination along elevation and azimuth (Dobreva et al., 2012)).

Each trial started with the presentation of the fixation dot for the duration of 1200 ms (Figure 1B). Next, participants were presented with dynamic Mondrians to one eye that suppressed their awareness of signals presented to the other eye (dCFS). After a random interval of 600-1100 ms, a sound was played from one of three potential locations. On the flash present trials, a white disc was presented in one of the four quadrants for 50 ms in synchrony with the sound. Response cues were presented 750 ms poststimulus. The Mondrian masks were presented on the screen until participants responded to all questions.

On each trial, participants responded to three questions in a self-paced manner within a total response window of 5 s: First, they reported the location of the beep (left, centre, right) via a three choice key press. Second, they rated the visibility of the flash (clear image, almost clear image, weak glimpse, not seen) according to a previously published Perceptual Awareness Scale (Ramsøy & Overgaard, 2004) (PAS) via a four choice key press. This Perceptual Awareness Scale and experimenter’s explicit instructions encouraged participants to categorize trials as invisible, only if they were ‘completely invisible’. Third, they reported the location of the flash (upper or lower hemifield) via a two choice key press. Critically, we designed orthogonal spatial auditory and visual tasks to minimize decisional biases between visual and auditory localization responses. In order to minimize response interference between responding to the set of three questions, we ensured that the responses mapped to distinct sets of buttons (i.e. 9 different buttons in total). The button/hand assignment and the order of questions were counterbalanced across participants (either: sound location – visibility – flash location or: flash location – visibility – sound location).

This visibility judgment provided a subjective awareness criterion. Critically, we adjusted the flash’s luminance in adaptive staircases individually for each participant, such that the flash was visible on only 40% of the trials. This allowed us to quantify multisensory interactions as indexed by spatial ventriloquism (i.e. audiovisual spatial bias) for flashes that were visible (i.e. pooled over ‘clear image’, ‘almost clear image’, ‘weak glimpse’) or invisible (i.e. ‘not seen’, subjective awareness criterion ).

### Experimental procedure

The study included a one day psychophysics experiment and four day EEG experiment (for the subset of 18 subjects).

Prior to all experiments, we adjusted the flash’s luminance in adaptive staircases (step size up: 8.8 cd/m^2^, step size down: 13.2 cd/m^2^), such that the flash was visible on 40% of the trials. The adaptive staircases were applied using a slightly modified experimental paradigm, in which the sound was presented always from the middle, the flash in one of the four quadrants and participants reported only the flash’s visibility using a binary choice (yes, no) and the flash’s location (up, down). After an initial long staircase (including at minimum 200 trials), we performed four times two interleaved adaptive staircases (convergence criterion: 8 reversals within the last 10 trials).

The one-day psychophysics experiment included a total of 8 experimental runs, resulting in a total of 432 trials (i.e. 64 trials for each flash present condition and 16 trials for each flash absent condition). In this initial psychophysics study the flash’s luminance was adjusted throughout the experiment with adaptive staircases to maintain a visibility level of approximately 40% (i.e. 60% of the trials were rated ‘not seen’, i.e. the lowest visibility based on the four level Perceptual Awareness Scale (Ramsøy & Overgaard, 2004; Sandberg et al., 2010)). To minimize the variability of the flash’s luminance during the psychophysics experiment we adjusted brightness of the flash in smaller step sizes (3.3 cd/m^2^) and only after 4 consecutive ‘not seen’ responses or after 3 consecutive ‘seen’ (including all three “partially visible” levels: clear, almost clear & weak glimpse) responses.

Importantly, for the EEG experiment we kept the flash’s luminance constant throughout the four-day-experiment based on the initial psychophysics experiment to ensure that differences in brain activity are related to stimulus perception (i.e. subjective flash visibility) rather than the physical properties of the flash. Each participant completed 76 experimental runs over 4 testing days, resulting in total in 4104 trials (i.e. 608 trials for each flash present condition and 152 trials for each flash absent condition).

### EEG data acquisition and pre-processing

Continuous EEG signals were recorded from 60 channels using Ag/AgCl active electrodes arranged in 10-20 layout (ActiCap, Brain Products GmbH, Gilching, Germany) at a sampling rate of 1000Hz (for technical reasons 6 sessions spread across 3 subjects were recorded at a sampling rate of 500Hz), referenced at FCz. Four electrodes were used for EOG recording (2 placed above and below right eye and two on the side of each eye). Impedances were kept below 10kΩ for EEG channels and 20kΩ for EOG channels. On each testing day 3D coordinates of EEG electrodes were digitized using Polhemus Fastrack (Polhemus Corp., Colchester, US).

Preprocessing was performed using MATLAB R2016b (Mathworks, Natick, Massachusetts) and Fieldtrip toolbox (Oostenveld et al., 2011). Muscle artefacts and noisy channels (0.8 channels on average) were identified based on visual inspection. Continuous EEG signals were high-pass filtered to 0.1 Hz and low-pass filtered to 30Hz. Independent component analysis (ICA) was applied to the non-noisy channels to correct for eye movements and heartbeat artefacts. Eye blink and heartbeat-related components were identified based on visual inspection of component topographies and time-courses. Between 2 and 6 ICA components were removed (3.1 components on average). Noisy channels were interpolated using weighted average of neighbouring channels, based on sensor positions from Polhemus recordings. Data were segmented into -0.15:0.7s epochs relative to target stimulus (i.e. flash + white noise burst stimulus) onset and re-referenced to average reference. After re-referencing, the FCz electrode was appended, so that signals from 61 channels were entered into the analysis. Trials containing artefacts were rejected. Furthermore, trials were rejected if they included eye blinks that overlapped in time with the presentation of the flash. The data were baseline corrected and downsampled to 60Hz.

#### Behavioural analysis for psychophysics and EEG experiment

The behavioural data of the initial psychophysics study and the EEG experiment are reported separately. All figures show only the results from the EEG experiment.

For data analysis, we reduced the four visibility levels to two visibility levels: 1. Visible = ‘clear image’, ‘almost clear image’ or ‘weak glimpse’ and 2. Invisible = ‘not seen’. Further, we pooled over flashes in upper and lower fields given their small elevation (i.e. ± 1.2° visual angle).

For each participant, we coded his/her sound location responses as -1 for left, 0 for centre and 1 for right across trials. We estimated the *perceived (or reported) sound location* for each of the 2 (flash location: left, right) x 3 (sound location: left, centre, right) conditions by averaging the localization responses across trials. Next, we averaged the perceived sound locations separately for trials on which the flash was presented on the left and right and computed the difference in average perceived sound location for ‘visual right’ minus ‘visual left’ trials as an index of the spatial ventriloquist effect. A positive value of this index indicates that subject’s perceived the sound location as shifted towards the visual stimulus location (i.e. attraction). A negative value indicates that it is shifted away from the visual stimulus location (i.e. repulsion). A ventriloquist effect of zero means that participants were not influenced consistently across trials by the location of the flash.

This difference in perceived sound location, i.e. the ventriloquist effect, was then used as the dependent variable for all subsequent analyses. If the visual signal location attracts the perceived sound location, we would expect the difference to be significantly greater than zero.

### Spatial ventriloquism for visible and invisible trials

We investigated whether the spatial ventriloquist effect was present independently for both invisible and visible flashes. Hence, we computed the ventriloquist effect separately for trials on which visual signals were judged visible (i.e. pooled over ‘clear image’, ‘almost clear image’, ‘weak glimpse’) or invisible (i.e. ‘not seen’) and tested whether the ventriloquist effect was significantly greater than zero in right-tailed one sample t-tests independently for visible and invisible trials.

Moreover, for the behavioural data from the four-day EEG experiment alone we also tested, whether the ventriloquist effect can be observed for invisible flashes, when participants mislocated the flash. This analysis was not possible for the psychophysics experiment in which we acquired data only in one single acquisition session, because the number of trials was not sufficient for this more refined sub-analysis.

### Correlation between visible and invisible ventriloquist effects

We investigated whether participants that show a strong (resp. weak) ventriloquist effect for invisible trials also exhibit a strong (resp. weak) ventriloquist effect for visible trials by testing for a Pearson correlation between the ventriloquist effects for visible and invisible flashes (across-trial mean for each observer) over observers. A significant correlation provides initial evidence that the neural mechanisms and circuitries underlying the ventriloquist effects for visible and invisible flashes may be shared.

### Influence of question order on flash localization accuracy and ventriloquism

As described earlier we counterbalanced the task order (either: sound location – visibility – flash location or: flash location – visibility – sound location) across participants to account for the fact that the order of the questions can influence observer’s performance accuracy on the different tasks. For instance, as a result of memory noise, the order of questions may influence observers’ flash localization accuracy and the size of the ventriloquist effect.

As a consequence, the question order could also influence the inclusion of participants. For instance, we may include a smaller number of subjects that were presented with the sound task first, because their flash localization accuracy for visible flashes may have been below our inclusion criterion. Conversely, we may include less participants with the flash task first, because their localization accuracy for invisible flashes may have been too high. We assessed whether the proportion of participants who had the flash localization first or last differed between the full sample of participants and the sample of participant who was included in the EEG experiment with a Chi-square tests.

In the EEG study, we also compared flash localization accuracy and the size of spatial ventriloquism between the groups of observers that performed the flash localization as first and last task using a two-sample t-test.

The behavioural data analysis was performed in MATLAB (Mathworks, Natick, Massachusetts), Bayes Factors were computed in JASP (Marsman & Wagenmakers, 2017). We report p-values and Bayes factors from two-sided tests unless otherwise stated.

#### EEG - Multivariate pattern analysis

Using multivariate pattern analysis, we addressed two questions: 1. To what extent is the location of the flash encoded in neural activity (i.e. classification accuracy based on EEG activity pattern)? 2. How does the location of the flash influence the neural encoding of the sound location (i.e. neural ventriloquist effect). Critically, we examined how flash and sound encoding depend on whether observers i. rated the flash as visible or invisible and ii. located the flash correctly or incorrectly. For sound encoding, we also compared trials with and without a ventriloquist illusion. We performed these analyses for the entire time from 0 to 650 ms post-stimulus and in a temporally-resolved fashion using sliding 50 ms time windows.

All multivariate analyses were performed in MATLAB (Mathworks, Natick, Massachusetts) using CoSMoMVPA toolbox (Oosterhof et al., 2016) and Libsvm package (Chang & Lin, 2011). EEG signals were z-normalized in each channel with normalization parameters from training set being applied to left out test set. In a leave one-fold out crossvalidation scheme (with 10 folds) we trained a support vector machine (with the C parameter set to 1) on single trial EEG activity patterns pertaining to i. the entire time window from 0 to 650 ms poststimulus (i.e. number of channels x 40 time samples, given a 60Hz sampling rate) or ii. overlapping 50 ms sliding time windows (i.e. number of channels x 3 time samples, given a 60Hz sampling rate).

### Decoding left vs. right flash location

Using support vector classification we investigated how the brain encodes flash location depending on observers’ visibility and flash localization accuracy. To ensure unbiased analyses we equated the number of trials of each of the audio-visual conditions (4 flash locations x 3 sound locations) in each of the training and testing folds. To further minimize confounding neural activity from sound processing we limited the analysis to 10 parieto-occipital EEG channels.

First, using a leave one-fold out crossvalidation scheme (with 10 folds) we trained a linear SVM to classify left vs. right flash locations (counterbalanced for up vs. down flash and sound locations) separately for flashes judged visible and invisible. Because the training data for left and right flash conditions were matched for the three sound locations, the EEG patterns that discriminate between left and right flashes were not confounded by sound location.

Second, we trained a SVM to classify left vs. right flash location on visible trials and then generalized this SVM to invisible trials i. irrespective of flash localization accuracy and ii. separately for trials on which observers located the flash correctly or incorrectly.

Flash decoding accuracy was entered into one sided permutation-tests to assess whether flash decoding accuracy was better than chance based on decoding from EEG activation pattern of the entire time window. For the sliding time window analysis we corrected for multiple comparisons across the entire [0 650] ms time window using the Threshold Free Cluster Enhancement procedure (Smith & Nichols, 2009) with one sided sign-permutation test (based on 10000 iterations) as implemented in CoSMoMVPA (Oosterhof et al., 2016). Unless stated otherwise, results are reported at p < 0.05 corrected for multiple comparisons across the specified time window.

### Decoding sound location and computing the neural ventriloquist effect

To investigate how the location of a flash influences the neural encoding of the location of a synchronous sound we trained a support vector regression (SVR) model to learn the mapping from EEG activity patterns to sound location in the external world based on unisensory auditory trials (left, middle, right). This training on auditory trials only ensures that the support vector regression model only uses EEG features related to sound processing. We used this trained SVR model to predict the sound location from EEG activity patterns of audiovisual trials. We computed a neural ventriloquist effect by subtracting the decoded sound location for flash right trials from flash left trials (i.e. as in our behavioural analysis). We assessed whether this ‘neural ventriloquist effect’ depends on flash visibility. For invisible trials only, we also assessed whether the neural ventriloquist effect depends on whether observers located the flash correctly or incorrectly. For visible trials, we could not perform this analysis, because observers did not mislocate a sufficient number of visible flashes.

Using one sided t-tests we investigated whether the neural ventriloquist effect was greater than zero based on decoding from EEG activation pattern of the entire time window separately for visible, invisible, invisible correctly localized and invisible incorrectly localized flashes. Further two sided paired t-tests were used to assess whether the neural ventriloquist effect differed between visible and invisible trials, as well as for invisible correct and invisible incorrect trials. For the sliding time window analysis we report p-values corrected for multiple comparisons using the Threshold Free Cluster Enhancement procedure ((Smith & Nichols, 2009), see above). Unless stated otherwise, results are reported at p < 0.05 corrected for multiple comparisons across the entire time window.

### Assessing the relationship between behavioural and neural ventriloquist

To assess the relationship between neural and behavioural ventriloquist effects we compared the neural ventriloquist effect for trials on which observers did or did not experience a perceptual ventriloquist illusion. Because the effect of behavioural ventriloquism on neural ventriloquism can be assessed in an unbiased fashion only for sound centre trials, this analysis was limited to those sound centre trials.

First, we assessed the neural ventriloquist effects for trials with vs. without ventriloquist illusion in separate paired-t-tests for visible, invisible, invisible correctly localized and invisible incorrectly localized flash trials.

Second, we entered the neural ventriloquist effect computed over the entire 0-650 ms time window into two repeated measures ANOVAs (n.b. we acknowledge that we transformed dependent variables [i.e. visibility, flash localization accuracy, behavioural ventriloquist effect] into independent factors):

The first 2(visibility: visible vs. invisible) x 2(behavioural ventriloquist effect: present vs. absent) repeated measures ANOVA assessed the statistical effect of observers’ visibility and behavioural ventriloquist effects on their neural ventriloquist effect. In the second 2 (flash localization accuracy: correct vs. incorrect) x 2(behavioural ventriloquist effect: present vs. absent) repeated measures ANOVA for the neural ventriloquist effect, we focussed on invisible trials only (n.b. there are no sufficient visible flash trials with incorrect localization.

The repeated measures ANOVAs and Bayesian statistics were computed in JASP (Marsman & Wagenmakers, 2017).

The neural ventriloquist effect (computed from the sound locations decoded from the entire time window or sliding time window) were entered into two sided t-tests. For the sliding time window analysis we corrected for multiple comparisons across the entire [0 650] ms time window.

### Influence of question order on decoding accuracy of flash location and neural ventriloquism

In a control analysis (as in our behavioural analysis) we assessed the influence of question order by comparing the decoding accuracy for flash location and the neural ventriloquist, computed over the entire window, between groups of subjects who performed flash or sound localization tasks first (i.e. to assess effects of question order).

## Notes

### Competing Interest Statement

The authors have declared no competing interest.

### Funding

- ERC: 101096659 ERC grant Making Sense, 309349 ERC grant MultSens

